# From sequences to therapeutics: Machine learning predicts chemically modified siRNA Activity

**DOI:** 10.1101/2023.08.16.553554

**Authors:** Dominic D. Martinelli

## Abstract

Small interfering RNAs (siRNAs) exemplify the promise of genetic medicine in the discovery of novel therapeutic modalities. Their ability to selectively suppress gene expression makes them ideal candidates for development as oligonucleotide pharmaceuticals. Recent advancements in machine learning (ML) have facilitated unmodified siRNA design and efficacy prediction, but a model trained to predict the silencing activity of siRNAs with diverse chemical modification patterns has yet to be published, despite the importance of such chemical modifications in designing siRNAs with the potential to advance to the clinic. This study presents the first application of ML to classify efficient chemically modified siRNAs from sequence and chemical modification patterns alone. Three algorithms are evaluated at three classification thresholds and compared according to sensitivity, specificity, consistency of feature weights with empirical knowledge, and performance on an external validation dataset. Finally, possible directions for future research are proposed.

## 1 Introduction

RNA interference (RNAi) is a natural biological mechanism that mediates specific gene silencing. RNAi has been leveraged not only as a research tool but also as a promising addition to genetic medicine’s contributions to the present diversity of therapeutic modalities [1]. One such class of RNA that mediates interference, siRNA, yields particularly specific gene suppression. making it an exceptional candidate for applications in drug discovery and development [2]. When Dicer, an enzyme of the RNAse III family, cleaves either double-stranded or hairpin RNA, siRNA duplexes are produced. These siRNAs are then bound by the RNA-induced silencing complex (RISC). One component of RISC, Argonaute-2, unwinds the siRNA such that the duplex is separated into single-stranded RNA. The siRNA can then align with its mRNA target, at which point it cuts the mRNA, inhibiting translation. A schematic of this mechanism is illustrated in Figure 1.

**Figure 1.**
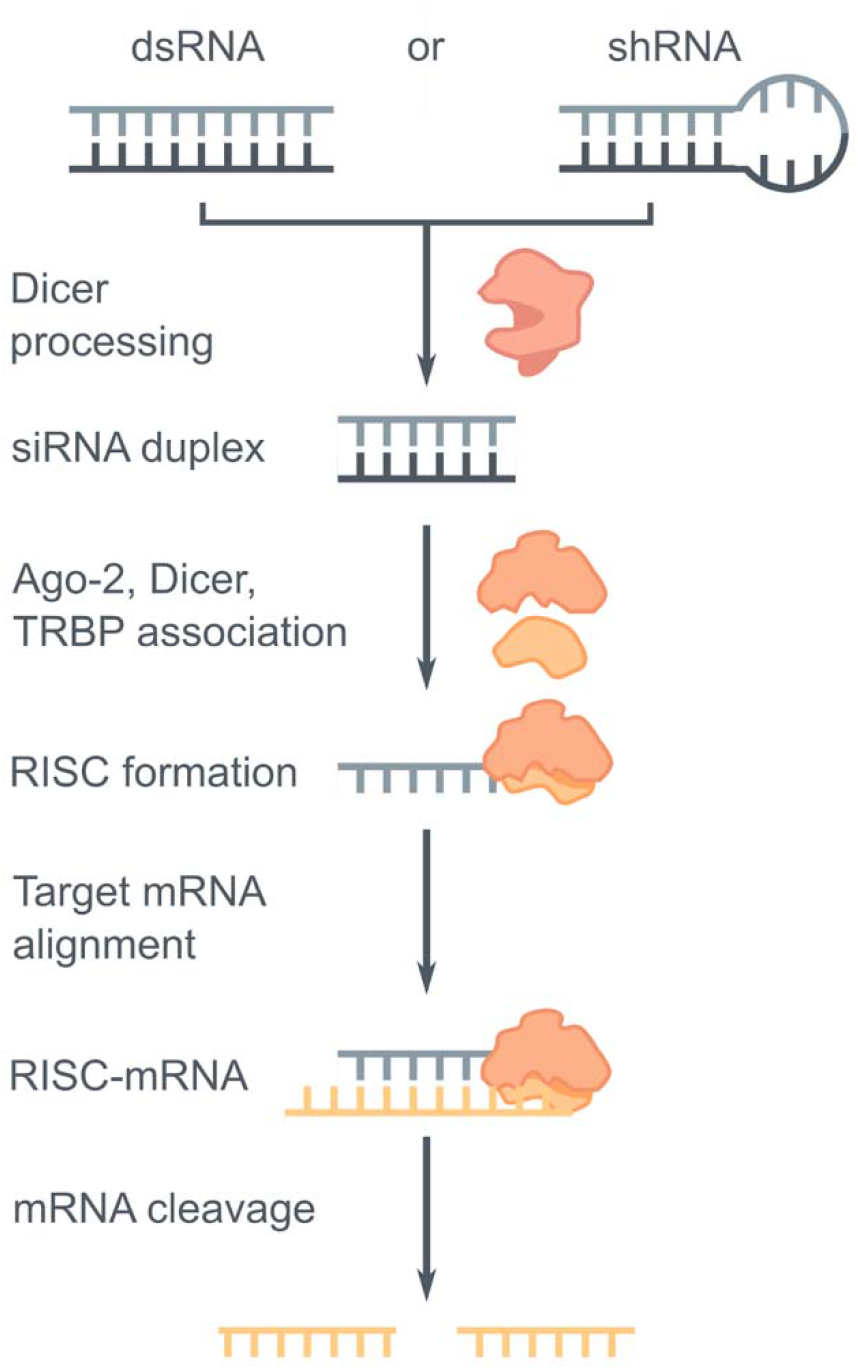
The mechanism of action of siRNA-mediated RNAi.

Chemical modifications are often introduced to the structure of siRNA to improve its developability [3]. This holds true for all FDA-approved siRNA therapeutics to date [4]. Such chemical modifications may reduce off-target effects, which is a major concern with any method of RNAi. Modifications can also enhance binding affinity while simultaneously improving stability where desirable. Several factors determine the tolerability of chemical modifications, including positional context, bulkiness, presence of other modifications or frequency of a single modification, nucleotide composition, and whether the modification is introduced to the sense or antisense strand. siRNA conjugate delivery platforms have also amassed popularity as an application of traditional medicinal chemistry techniques to this novel therapeutic modality [5].

ML, a subtype of artificial intelligence, has been applied to a variety of tasks in the design of nucleotide-based therapeutics, including siRNAs. Early publications apply relatively simple algorithms to this prediction task, yielding encouraging results [6]–[8]. The first application of an artificial neural network (ANN) to the curation of a library of therapeutic siRNAs produced one of the first large datasets in the field, facilitating further research on ML for therapeutic oligonucleotide design [9]. However, an algorithm that predicts the relationship between siRNA chemical modification patterns and efficiency has yet to be published. This area is especially important to address in the context of gene therapy design, as all FDA-approved siRNA therapeutics to date are chemically modified [4]. Hence, a model trained to predict silencing activity only for unmodified siRNA sequences may possess minimal relevance to a drug development context. It is therefore imperative to determine whether ML approaches to siRNA efficacy prediction could extend beyond the narrow applicability domain of unmodified siRNAs.

In this article, three ML techniques are trained on a dataset of chemically modified siRNAs to predict their silencing efficiency from sequence and chemical modification patterns. These models are assessed with respect to a variety of factors, including specificity, sensitivity, predictive uncertainty, and interpretability. Weights assigned to positionspecific chemical modifications are interpreted and discussed in the context of empirical siRNA design principles. The constructed models are then rigorously evaluated against an external validation dataset. This publication represents the first application of ML to understand the relationship between siRNA sequence, chemical modification pattern, and silencing efficacy. It serves as a proof-of-concept foundational study for future research on data-driven approaches to chemically modified therapeutic siRNA design, especially in silico techniques.

## 2 Methods

### 2.1 Dataset curation

The training and testing datasets were derived from the siRNAmod database, an archive of chemically-modified and control siRNA sequences reported in the literature, along with their respective biological activities [10]. The validation set, which is made accessible in the Supplementary Information section, was manually collected by extracting siRNA sequences and their corresponding inhibitory activities from published articles not included in the siRNAmod database. It comprises a total of 57 siRNAs, 53 of which are chemically modified, sourced from six documents. The PubMed IDs of these documents are specified in the Supplementary Information. To ensure sufficient data quantity and quality, included siRNAs were subject to the following criteria:

- Length of 21 nucleotide bases;
- The exact biological inhibition percentage corresponding to the siRNA is reported in the database;
- If chemical modifications are present, the specified modifications must exist at least 10 times within the dataset.

These criteria were applied to facilitate the process of learning from the data. It may be possible to use ML to predict the effectiveness of chemical modification patterns of excluded chemical modifications as more data is published. Despite the exclusivity inherent to any data criteria, the standards established for this study do not come at the cost of sample diversity. Seven common chemical modifications, depicted in Figure 2, were represented in the dataset. The abundance of each modification in the dataset and the distribution of efficiency values will be presented with the results of the experiment in Section 3.1. Moreover, the training, testing, and validation datasets are included as Supplementary Information.

**Figure 2.**
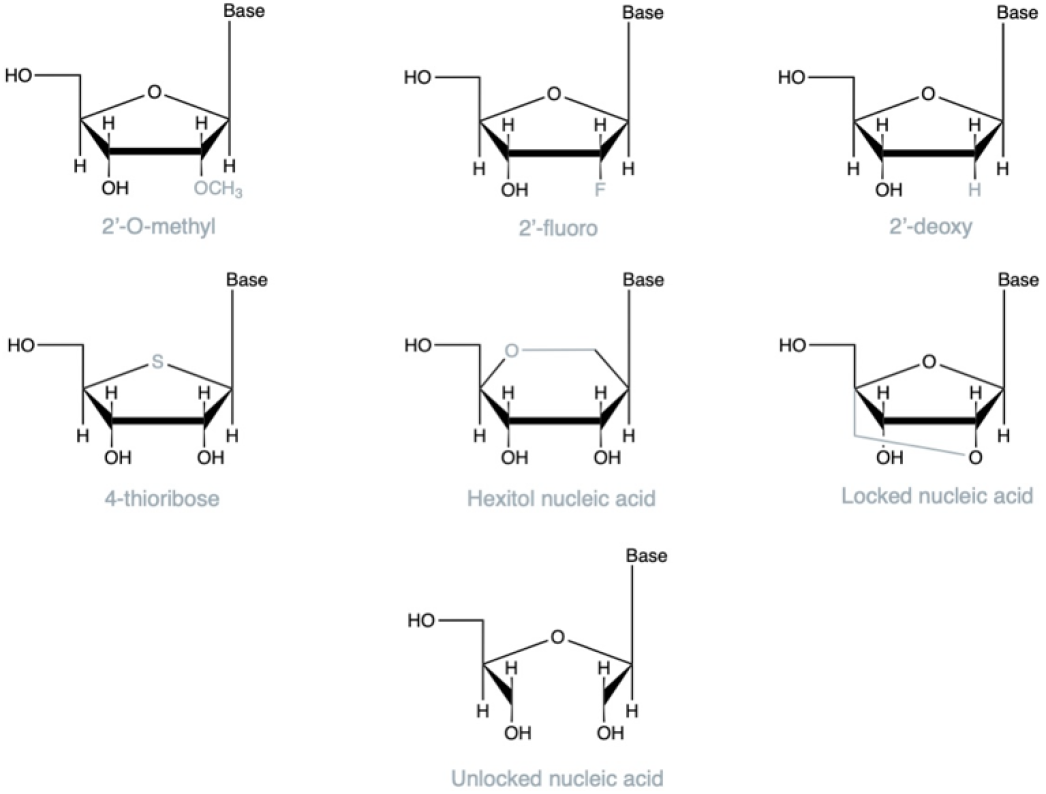
The ribose chemical modifications represented in the dataset.

### 2.2 Efficiency classification thresholds

While a knockdown efficiency of 70% typically suffices for effective gene silencing [11]–[13], a more dramatic reduction in gene expression is sometimes sought. To verify robustness to multiple definitions of inhibitory efficiency, three classification thresholds—60%, 70%, and 80% biological inhibition—were applied to each sample. Classifying siRNAs based on a higher threshold would not yield predictive models due to the dramatic class imbalance that would be present in such a model. This assessment of performance consistency at different thresholds was also employed to identify potential discrepancies in model performance when trained on datasets with different minimum efficiency values.

### 2.3 Featurization

To limit the computational cost of the problem without sacrificing performance, features were limited to the base sequence and chemical modification pattern of each sample. One-hot encoded arrays were generated for each of these two broad feature categories from both the sense and antisense strands. These arrays were then concatenated along the y-axis, producing a complete encoded feature vector with a length of 546, representing each base in the 21-nt long sense sequence, the antisense sequence of the same length, and any combination of chemical modifications at each position along either siRNA strand.

### 2.4 Model construction

The Python package scikit-learn [14] was utilized to split the data into training and testing sets, build the models assessed in this study, evaluate model performance, and perform cross-validation. 75% of the dataset was allocated to training the model, while 25% was conserved for testing. A random seed of 42 was specified at all classification thresholds, facilitating reproducibility and consistency. scikit-learn’s train_test_split function distributes positive and negative samples evenly while minimizing bias, making it ideal for fair model evaluation.

Three ML classifiers—a random forest, a multi-layer perceptron ANN, and a linear support vector machine (SVM)—were trained and tested on the initial dataset at each of the two defined efficiency thresholds. These models were selected because of their well-established quality performance as predictors of unmodified siRNA efficiency [9], [15]–[17]. Moreover, they have diverse strengths and pitfalls, some of which are outlined in Table 1. The benefit of studying a collection of diverse models, rather than attempting to optimize a single model architecture, lies in the insight such studies can provide about the suitability of certain approaches in modeling novel data.

**Table 1.** Advantages and disadvantages of the three ML algorithms applied in this study.

| Model | Advantages | Disadvantages |
| --- | --- | --- |
| Random forest | Adapts to sparse data, learns diverse label encodings, handles high dimensionality, captures nonlinear relationships [18] | Some risk of overfitting, difficulty handling redundant or irrelevant features [19] |
| Multi-layer perceptron ANN | Able to capture the nonlinear relationships that often characterize biological activity data, computationally efficient, learn implicit relationships especially well [20] | Inferior interpretability, may introduce unnecessary computational complexity that risks overfitting, demands large set of training data [21] |
| SVM | Operates well in low-data settings, predictive of relationship between sparse data and target variables, minimal generalization error [22] | Can be difficult to interpret [23] |

The scikit-learn implementations of these models are relatively interpretable and are hence well-adapted to the research questions this article intends to address. For all models, the maximum number of allowed iterations was set to 5,000. This allowed all models to converge successfully. The random state was set to 42, ensuring reproducibility. Two hidden layers, with sizes of 62 and 12, respectively, were set for the multi-layer perceptron. The hidden layer sizes were decreased from the default value of 100 due to the relatively small, but still appropriate, size of the dataset. 1,000 estimators were integrated into the random forest model. All other parameters were set to their default values.

### 2.5 Performance evaluation and validation

The models were ranked according to the receiver operating characteristic (ROC) and precision-recall curve (PRC) area under the curve (AUC) values. The ROC metric considers the rates of true and false positives. Formulas 1 and 2 demonstrate the method of calculating true and false positive rates, respectively.

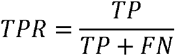

Formula 1. Calculation of the true positive rate (TPR) from binary classifier output.

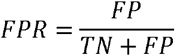

Formula 2. Calculation of the false positive rate (FPR) from binary classifier output.

The ROC AUC itself is computed according to Formula 3. This value is positively correlated with model performance.

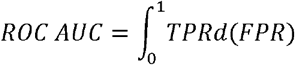

Formula 3. Calculation of the ROC AUC.

ROC AUC was preferred to accuracy as a metric of model performance for several reasons. Firstly, the data used to train this model was not uniformly distributed at any classification threshold, making accuracy less than ideal as a metric of model performance [24]. It is unlikely that, in a drug discovery setting, there will exist a perfect balance between positive and negative samples, so making this assumption would be inappropriate [25]. In addition, identifying the risk of false positives was deemed especially important given that such optimistic predictions directly contribute to the substantial cost of failure in drug development [26], [27]. While excluding siRNAs with the potential to succeed is also undesirable, it does not come with the same cost burden as excessively optimistic predictions. This trade-off is reasonable, given that discovering a single highly efficacious and developable siRNA is more meaningful in applied pharmaceutical research than designing a library of all possible sequences. To extract insights about the data features and relationships between chemical modification patterns and efficiency, weights were extracted from the highest-performing model.

The PRC was also used to evaluate the models. There is some concern that the ROC is overly optimistic when trained to classify imbalanced data, but the AUPRC may be more suitable for such datasets [28]. Hence, the AUPRC was computed to substantiate any conclusions drawn from evaluation of the ROC AUC. Formulas 4 and 5 demonstrate the method of calculating precision and recall, respectively.

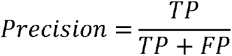

Formula 4. Calculation of precision from binary classifier output.

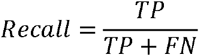

Formula 5. Calculation of recall from binary classifier output.

To more rigorously evaluate the highest-performing model, cross-validation was accomplished using the K-fold method [29]. A fold number of five was set for validation. Effective and ineffective siRNAs were evenly distributed to ensure fair comparison across folds. K-fold validation was performed with both the testing dataset and the manually curated validation dataset. The complete ML workflow executed in this study is illustrated in Figure 3.

**Figure 3.**
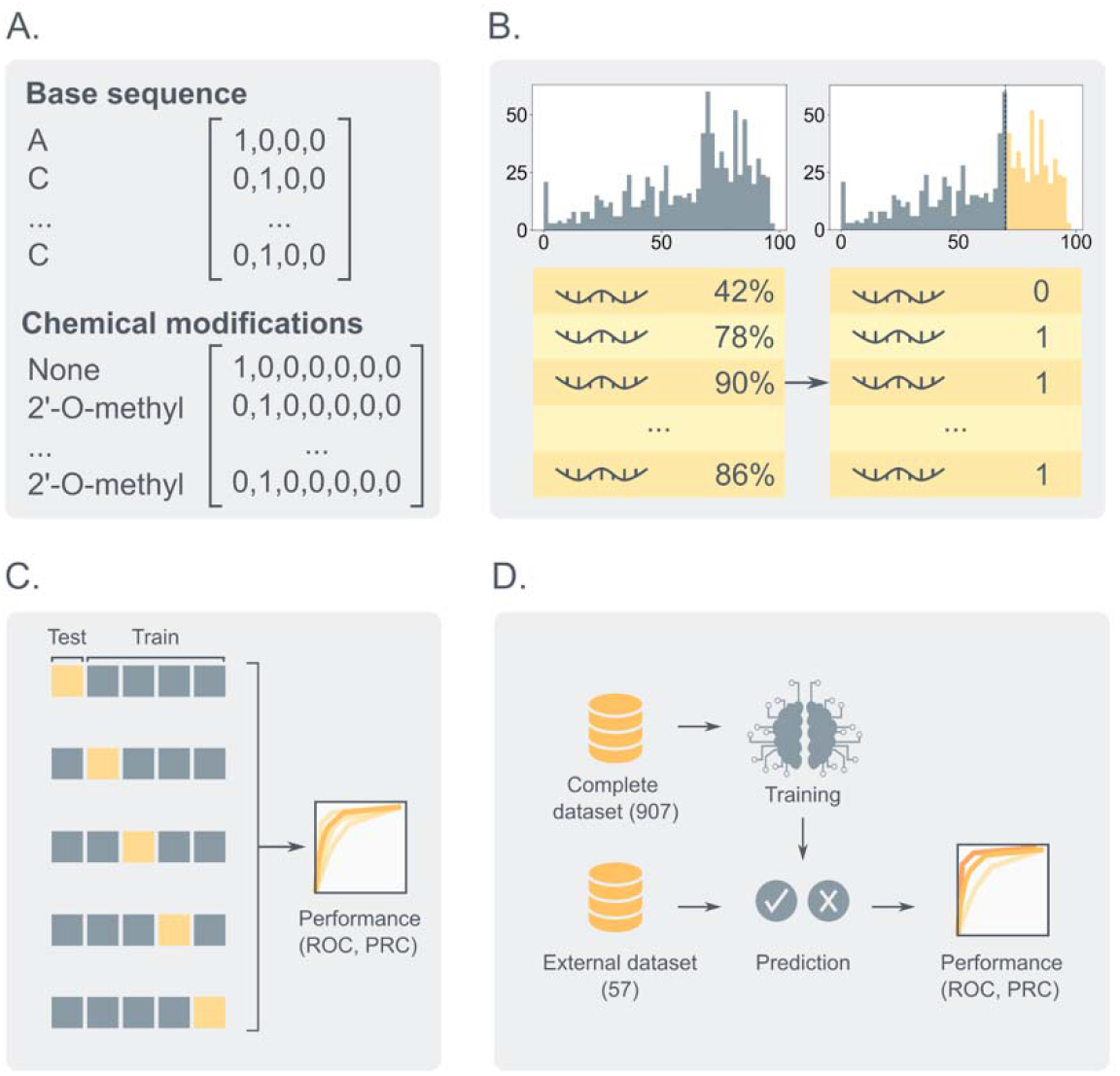
The procedure used to train, assess, and validate the ML models in this study. (A) Bases and chemical modifications are first one-hot encoded, then (B) labeled as efficacious (1) or inefficacious (0) according to the selected threshold. (C) Five-fold cross-validation is performed for each of the three models. Finally, (D) the trained models are evaluated on externally-sourced data.

## 3 Results and discussion

### 3.0.1 siRNA activity dataset

The protocol implemented to collect the dataset of 21-mer siRNAs, including chemically modified sequences with recorded activities, is described in the Methods section. It contains a total of 907 siRNAs, all of which are chemically modified. These siRNA records were extracted from 30 unique documents published either as patents or as scientific articles. In Figure 4, the distribution of the biological inhibition percentages across the siRNA dataset is visualized using a histogram. A diverse range of bioactivity values are represented in the dataset, although some reporting bias is reflected by the data’s moderate left skew.

**Figure 4.**
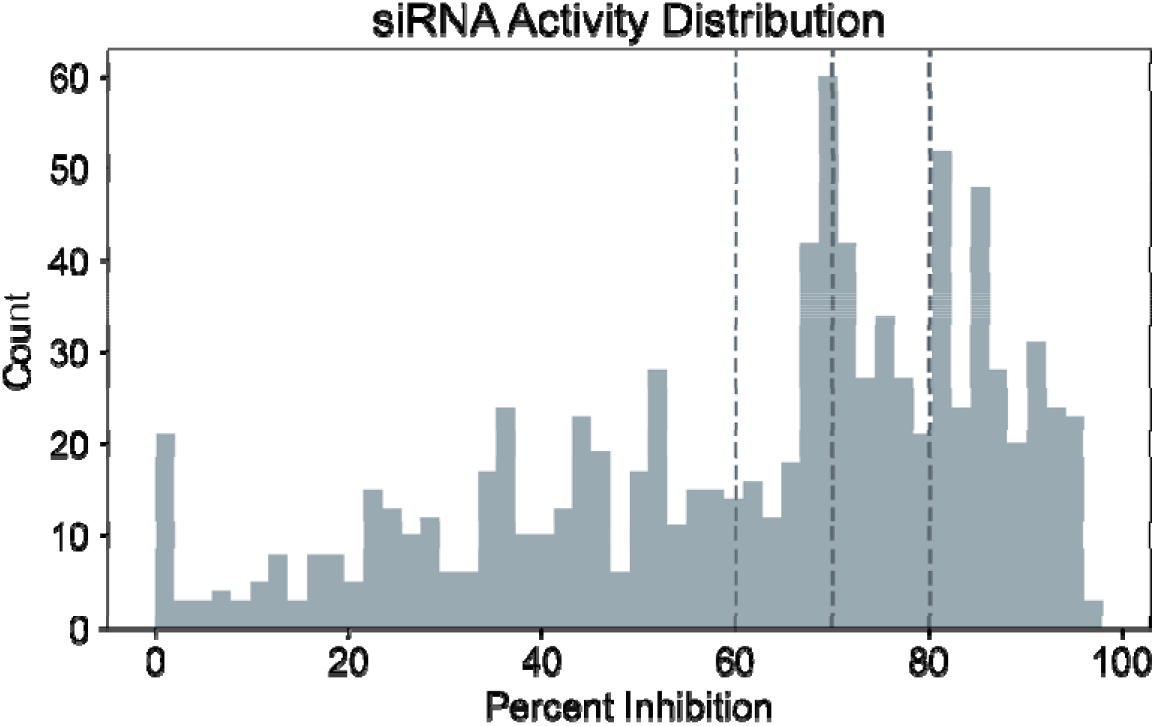
The distribution of activities represented in the dataset. Dashed lines represent the 60%, 70%, and 80% classification thresholds.

The frequencies of the types of chemical modifications used to characterize the siRNAs in this dataset are displayed in Figure 5. The relative prevalences of each chemical modification integrated into this model generally reflect their frequencies in the literature, with 2’-O-methyl and 2’-F modifications being the most common [10]. In addition to chemical modification patterns, positional base identities were encoded to characterize individual siRNAs. Sequencelevel features are highly predictive of the biological activity of non-modified siRNAs [30], [31] and are trivial to compute, making them well-adapted for the purpose of ML. Although other parameters, such as thermodynamic properties or off-target activities, may enhance model performance [32], [33], they are not as readily available as sequence and modification patterns. This model is designed to operate even in low data settings, facilitating application to novel chemically modified siRNA sequences.

**Figure 5.**
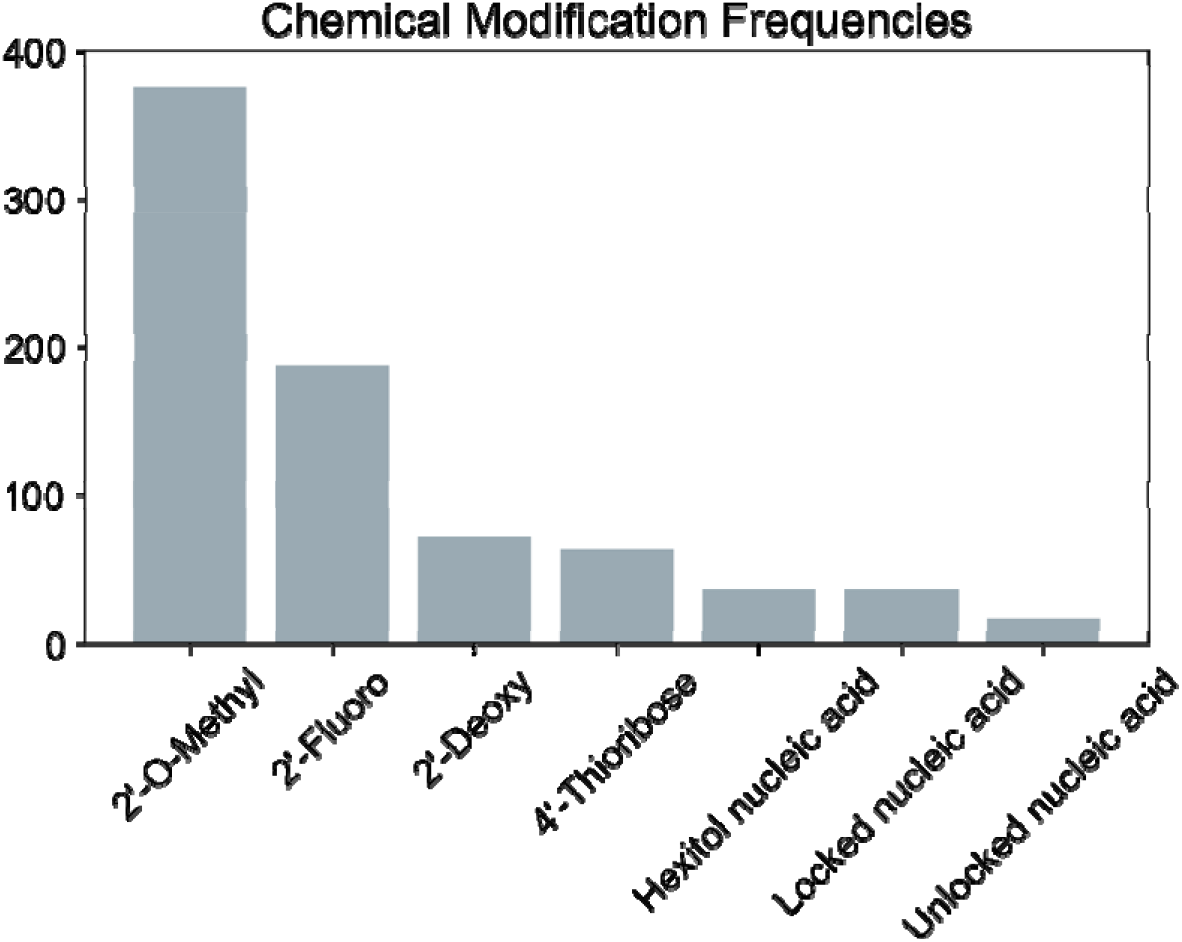
The appearance frequencies of the chemical modifications in the dataset.

In the literature, various thresholds for efficient siRNA silencing are defined. This threshold varies across publications, explicating the need for a model with robust predictive performance at multiple levels of inhibitory activity. As described in the Methods section, three activity thresholds–60%, 70%, and 80%–were utilized to classify efficacious siRNAs.

### 3.0.2 Machine learning models are robust to multiple class thresholds

According to both the ROC AUC and AUPRC metrics, performance was consistently high for all class definitions. For all models and at all thresholds, the average ROC AUC and AUPRC values invariably exceeded 0.70. Over the five K-folds for each of the three thresholds, the minimum and maximum ROC AUC scores were 0.62 and 0.95, respectively. The lowest and highest AUCPRs were 0.51 and 0.95, respectively. This demonstrates that the predictive ability of ML models extends beyond unmodified siRNAs to chemically modified sequences. The cross-validated performance of the random forest, multilayer perceptron, and SVM classifiers at each threshold are displayed in Figure 6.

**Figure 6.**
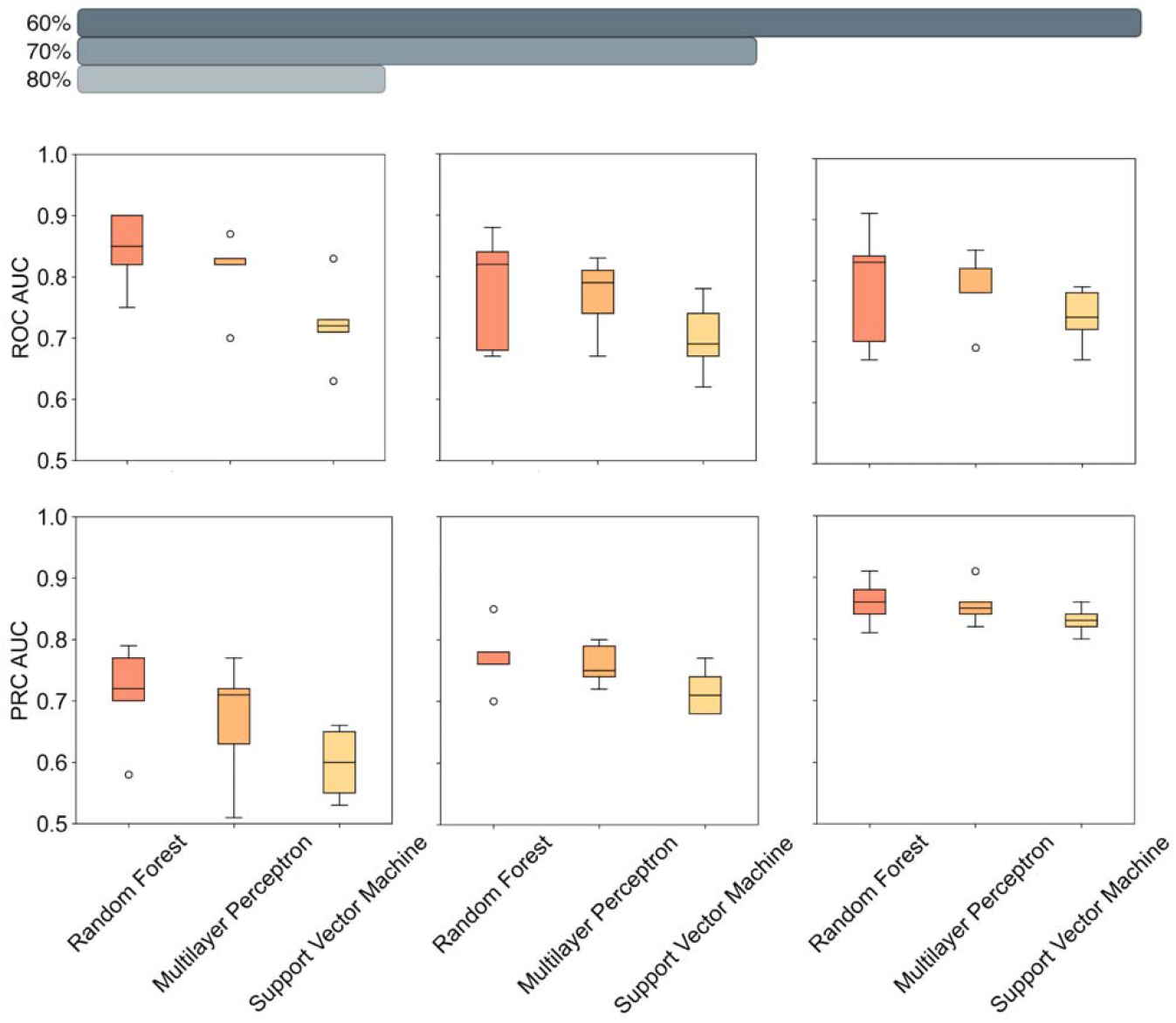
The results of the five-fold cross-validation for each model at three thresholds.

### 3.0.3 Random forests outperform non-ensemble models in predicting chemically modified siRNA efficiency

The average performance of the random forest algorithm, as measured by the median ROC AUC across five K-folds, exceeded or matched that of the multilayer perceptron and support vector machine at all thresholds. This is consistent with earlier findings demonstrating the superior performance of random forests in learning relationships between siRNA sequence and efficiency relative to techniques that leverage only a single predictor [34]. However, the superiority of the multilayer perceptron relative to the support vector machine was surprising, as prior comparative studies demonstrate that support vector machines surpass artificial neural networks when trained to predict biological activity from unmodified siRNA data [17]. Several variables, including noise, the number of features, and the nature of the prediction task (i.e., binary classification versus regression), might contribute to this discrepancy. Additionally, the superior performances of the RF and MLP ANN come at the cost of interpretability. To more precisely demonstrate variability and confidence in predictions from the best-performing model, ROCs and PRCs for each instance of cross-validation at the studied thresholds for the random forest are plotted in Figure 7.

**Figure 7.**
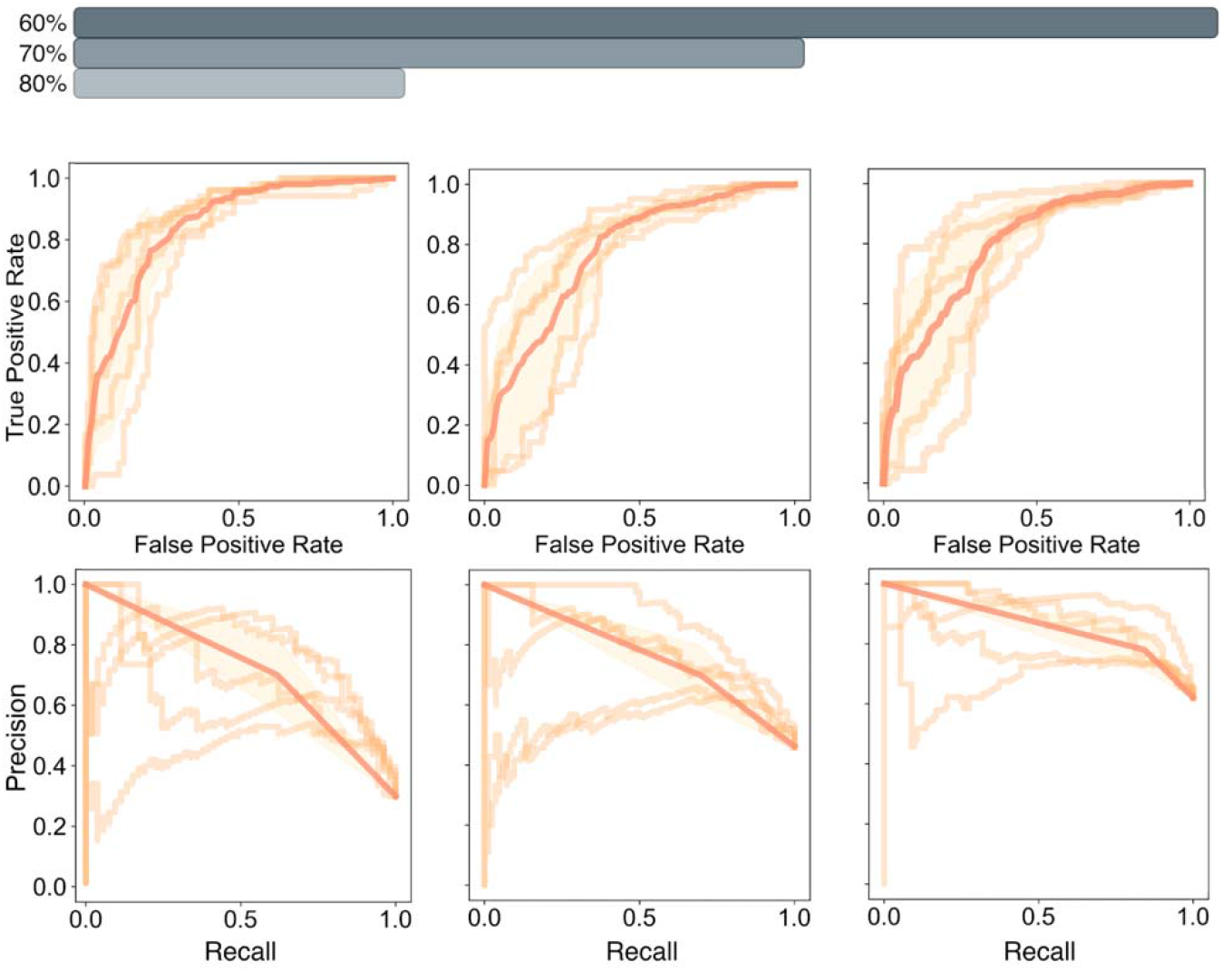
The cross-validated performance of the random forest classifier.

### 3.0.4 Learned feature importances reflect empirical knowledge and bias

As the feature weights assigned by the random forest model at the 60% classification threshold (Figure 8) substantiate, moderately stabilizing modifications (e.g., single-position 2’-O-methyl, LNA, and 2’-F substitutions) at the 5’ end of the sense strand enhance the activity of the antisense strand [35]. Likewise, destabilization of the 3’ end of the sense strand tends to augment antisense silencing efficacy. This effect is also important, as can be deduced from the relatively high weight assigned to the destabilizing 2’-deoxy modification at positions 20 and 21 of the sense strand. Additionally, the importance of the 2’-O-methyl modification at position 2 in the antisense strand, as reported by Jackson et al., seems to have been learned by the model [36].

**Figure 8.**
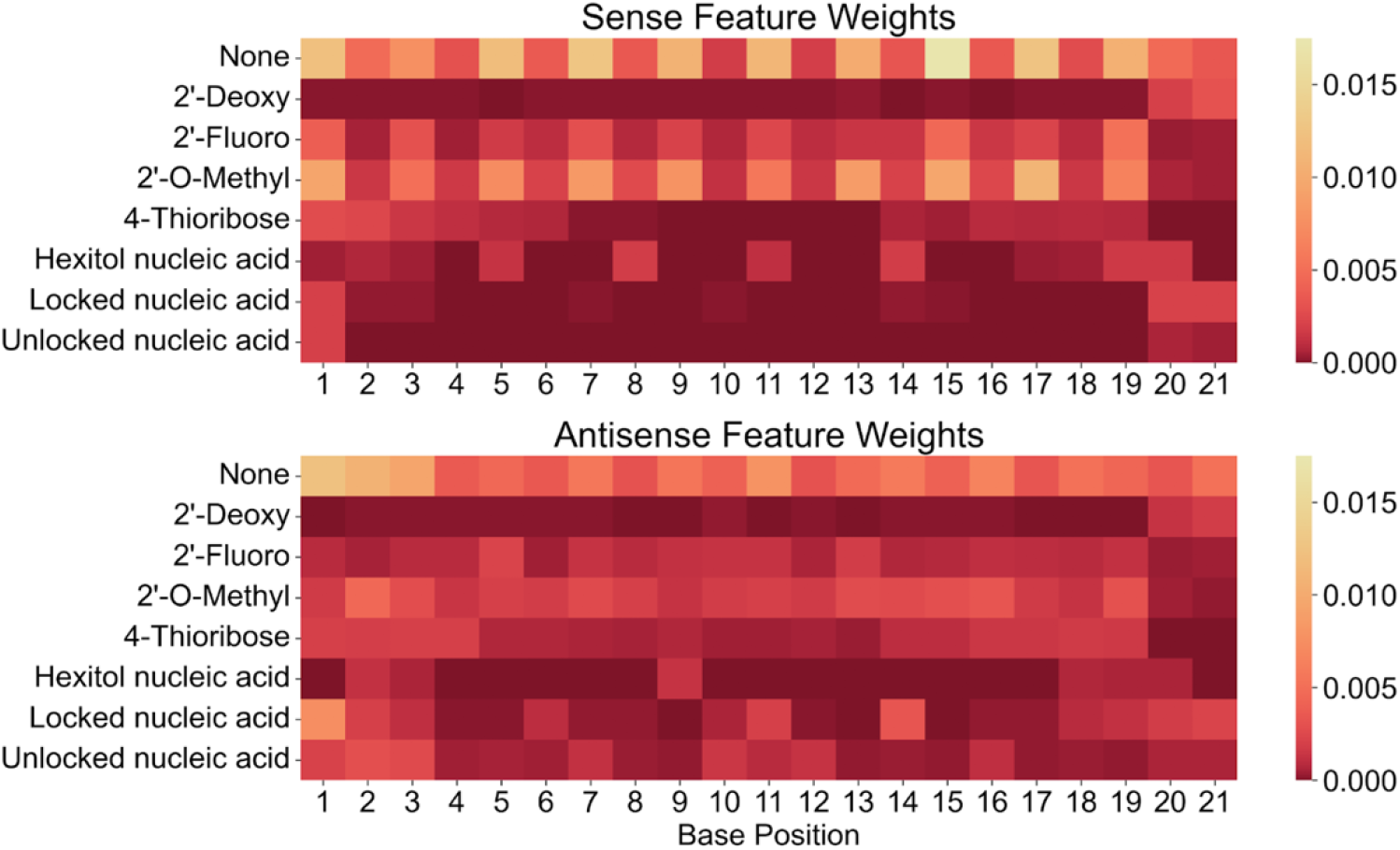
Weights assigned by the random forest classifier at the 60% classification threshold trained on the full dataset.

One challenge in interpreting the weights extracted using the precompiled random forest model is the ambiguity in the exact relationships learned by the model. For example, the importance of the LNA modification at the 5’ and 3’ ends of the antisense and sense strands is reported in the literature with conflicting data on its favorability [37]. Although the presence of this modification is clearly important to the RF classifier, it is unclear whether there exists a positive or negative association between the LNA modification and efficacy, as the feature importances exclusively comprise positive values.

A chemical modification pattern negatively associated with siRNA activity is the introduction of multiple stabilizing modifications [35]. For instance, the higher weight assigned to the 2’-O-methyl modification at position 1 may not reflect the importance of stability at that position specifically, but the fact that the presence of 2’-O-methyl at this position often correlates with the presence of other stabilizing modifications at the 5’ end of the sense strand. This demonstrates that familiarity with the data and its limitations is a prerequisite to effective interpretation of assigned weights. Moreover, some position-modification pairs are only representative of a single base identity-modification pair. Therefore, the model may not generalize to unconventional base identity-modification combinations. Because the feature weights assigned to base identities at each position in the sense and antisense strands were also recorded, however, identifying potential collinearity is straightforward.

### 3.0.5 Performance on validation dataset

The three models used in this study were retrained on the full set of chemically modified siRNAs from the siRNAmod database before being assessed on the validation set at the three specified classification thresholds. At the 70% classification threshold, all models attained a high ROC AUC value, indicating that the learned relationships between siRNA inhibitory potency, base sequence, and chemical modification pattern extrapolate to unseen experimental conditions. However, model performance on the validation dataset was inconsistent across thresholds, as was observed in the cross-validation assessment on the original dataset. The random forest was the most robust to threshold variation, while the ANN suffered the most from the variation. The superior performance at lower classification thresholds demonstrates the importance of balanced training data. There are far fewer siRNAs with a silencing efficacy at or above the 80% threshold, making the identification of such sequences difficult with a data-dependent approach. The amplitude of siRNA potency does not determine the triviality of classifying effective sequences, but it is interesting to note that some of the tested models were less performative at the 60% threshold, despite the improvement in class balance. Figure 9 displays the ROCs and PRCs for each model, along with their corresponding AUC values, facilitating graph interpretation.

**Figure 9.**
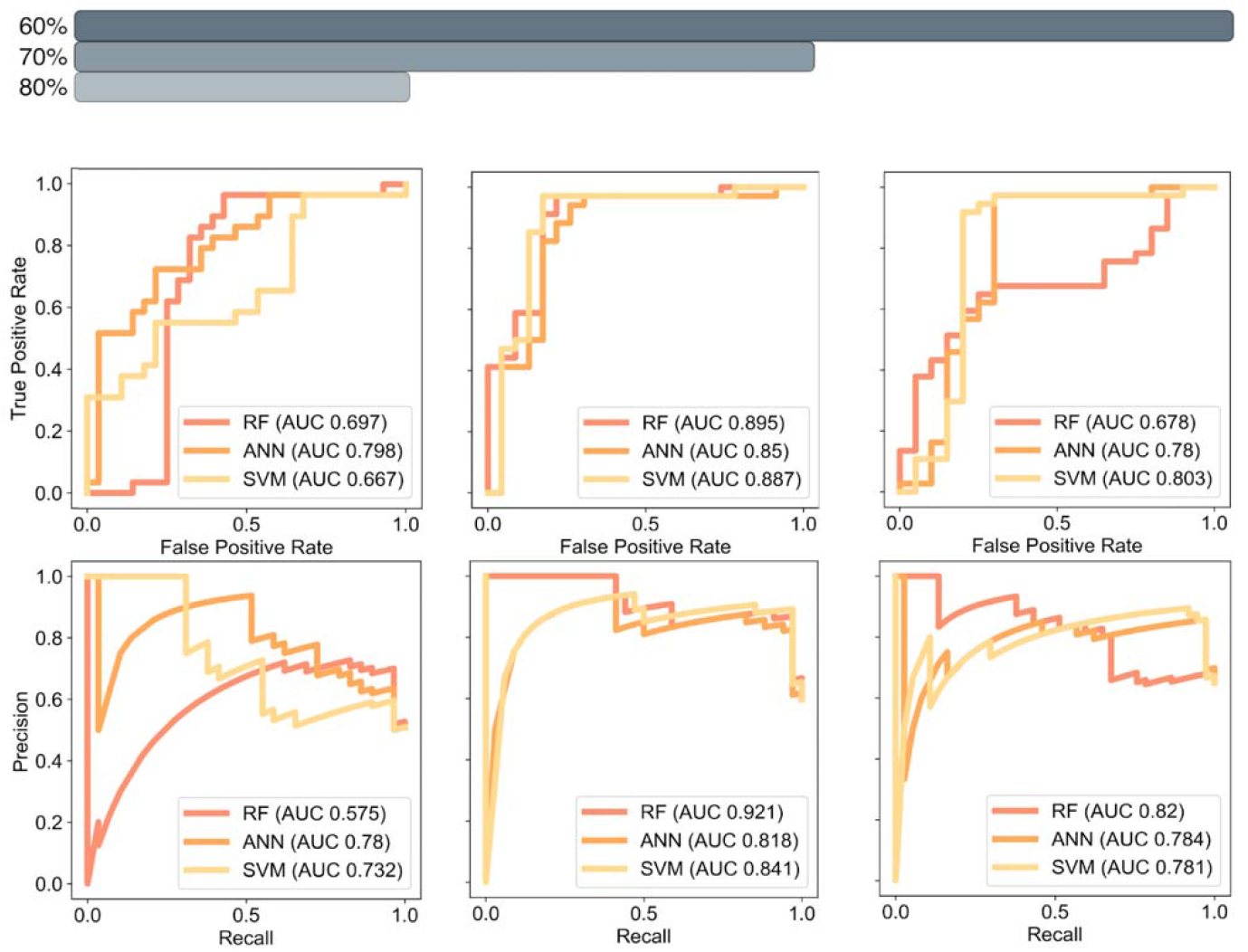
ROC and PR curves for the models validated with the external dataset.

### 3.0.6 Opportunities for future work

The data presented in this manuscript provide a robust foundation for further research, proving that ML models are able to learn to predict the efficiency of chemically modified siRNAs. As more data is published, it may be possible to predict the effect of novel chemical modifications, for example, by encoding information about the chemical structures of modifications rather than featurizing them with one-hot encoding. While such a featurization method satisfies the purpose of this proof-of-concept study, the flexibility afforded by the use of cheminformatics conventions would enable training on unseen modification types. Ebalunode and Zheng have already demonstrated that such descriptors can be used to represent the standard siRNA bases [38], but an investigation of their applied effectiveness in the modified oligonucleotide space has yet to be published. Hence, this report substantiates the hypothesis that extending the language of oligonucleotide chemistry to be learned using AI techniques is both possible and well within reach. Another area for future development concerns the construction of models that can predict the properties of siRNA sequences of variable length. The limitation of the datasets to oligonucleotides of 21 bases was a choice of convenience and does not imply that siRNAs with different lengths would be any more or less challenging to predict. Therefore, a model that is able to learn from data with variable dimensions would be a valuable elaboration on the methods proposed in this article.

## 4 Conclusions

In this study, ML is applied for the first time to predict the efficacies of chemically modified siRNAs with diverse modification patterns. Three well-established model architectures are trained, cross-validated, and assessed on an independent validation dataset to determine whether predictive models could learn from chemical modification patterns and base sequences. The results are promising, demonstrating the applicability of ML models to this prediction task. The ensemble method, an RF classifier, outperformed single-algorithm ML architectures, supporting earlier findings from studies on the use of ML to predict the efficacy of unmodified siRNAs. Features are not prioritized uniformly across models. The SVM classifier provided superior interpretability, despite its lower ROC AUC. Moreover, the constructed models retain predictive accuracy on validation data sourced externally, demonstrating that the learned feature weights reflect the rational basis for successful siRNA design, not experimental environment-exclusive sequence or chemical modification preferences. Further research may explore other methods of representing chemical modifications computationally and extending models to adapt to sequences of variable lengths.

## 5 Declaration of interests

The author declares no competing financial interests or personal relationships that could have appeared to influence the work reported in this paper.

## 6 Acknowledgements

The author thanks Ian Watson for constructive feedback and mentorship.

## Notes

### Competing Interest Statement

The authors have declared no competing interest.

